# Conditional control of universal CAR T cells by cleavable OFF-switch adaptors

**DOI:** 10.1101/2023.05.22.541664

**Authors:** Michael Kvorjak, Elisa Ruffo, Yaniv Tivon, Victor So, Avani B. Parikh, Alexander Deiters, Jason Lohmueller

## Abstract

As living drugs, engineered T cell therapies are revolutionizing disease treatment with their unique functional capabilities. However, they suffer from limitations of potentially unpredictable behavior, toxicities, and non-traditional pharmacokinetics. Engineering conditional control mechanisms responsive to tractable stimuli such as small molecules or light is thus highly desirable. We and others previously developed “universal” chimeric antigen receptors (CARs) that interact with co-administered antibody adaptors to direct target cell killing and T cell activation. Universal CARs are of high therapeutic interest due to their ability to simultaneously target multiple antigens on the same disease or different diseases by combining with adaptors to different antigens. Here, we further enhance the programmability and potential safety of universal CAR T cells by engineering OFF-switch adaptors that can conditionally control CAR activity, including T cell activation, target cell lysis, and transgene expression, in response to a small molecule or light stimulus. Moreover, in adaptor combination assays, OFF-switch adaptors were capable of orthogonal conditional targeting of multiple antigens simultaneously following Boolean logic. OFF-switch adaptors represent a robust new approach for precision targeting of universal CAR T cells with potential for enhanced safety.

## Introduction

Chimeric antigen receptors (CARs) are synthetic T cell receptors that specifically bind to antigens on the surface of target cells, leading to T cell activation and target cell lysis^1^. CARs consist of an extracellular antigen-binding domain fused to an extracellular spacer, transmembrane domain, and intracellular signaling domains. CAR T cell therapy, for which patients’ T cells are genetically engineered to express the CAR, has transformed the treatment of hematological cancers and shows tremendous promise for treating other diseases ^2, 3^. CAR T cell therapies targeting CD19 and BCMA positive hematological cancers are now FDA-approved and show remarkable therapeutic response rates, including durable cures in a fraction of patients^4^. Many new CAR T cell therapeutics are in development, targeting other antigens and diseases^5^.

Despite current success, CAR T cell therapy faces several major challenges to fulfilling its promise. Challenges include toxicities resulting from overactivity of CAR T cells (e.g., cytokine storm and neurotoxicity) and ON-target/OFF-disease activation that occurs when the target antigen is also expressed on non-diseased cells (e.g., FDA-approved anti-CD19 CAR T cell therapy causes B cell aplasia)^6^. Additionally, if the target antigen is lost, cancers can evade CAR T cell killing, resulting in disease relapse^7^. These limitations are of paramount concern with current therapies, as well as new therapies in development, as they ultimately lead to therapeutic failure^8, 9^.

To overcome some of these challenges, we and other groups, have engineered universal CARs that offer enhanced control over T cell function compared to traditional CAR T cells^3, 10, 11^. Instead of directly binding the target antigen, the CAR binds to a co-administered antibody adaptor specific to the target antigen. Binding of the CAR to the adaptor on the target cell surface leads to CAR T cell activation and target cell killing. This approach has several advantages over traditional CAR T cell therapies, including tuning CAR activity by dosing the adaptor molecule, changing targeting specificity by switching the adaptor to target multiple cancers with the same CAR T cell product, and avoiding tumor relapse by simultaneously targeting multiple antigens with co-administered adaptors^11^. We previously generated a universal CAR with the affinity-enhanced monomeric streptavidin (mSA2) protein as the adaptor binding domain^12^. This CAR interacts with high affinity to biotin-tagged adaptors to mediate T cell effector functions (Figure 1A). Several additional universal CAR technologies with different adaptor tag designs have been generated, and some are now being tested in clinical trials^13^. Despite the additional control that these technologies provide, safety concerns are still not fully addressed. The antigen specificity and ON-target/OFF-disease safety of universal CAR T cells are still a function of the adaptor binding profile. Even with extensive pre-clinical testing, toxicities from targeting new antigens are difficult to predict and can occur rapidly in patients^9, 14^. Furthermore, current OFF-switch approaches require turning off all CAR activity and are not tailored for specific switching of distinct antigens or in specific locations. Thus, additional means to switch OFF CAR activity would be desirable.

**Figure 1.**
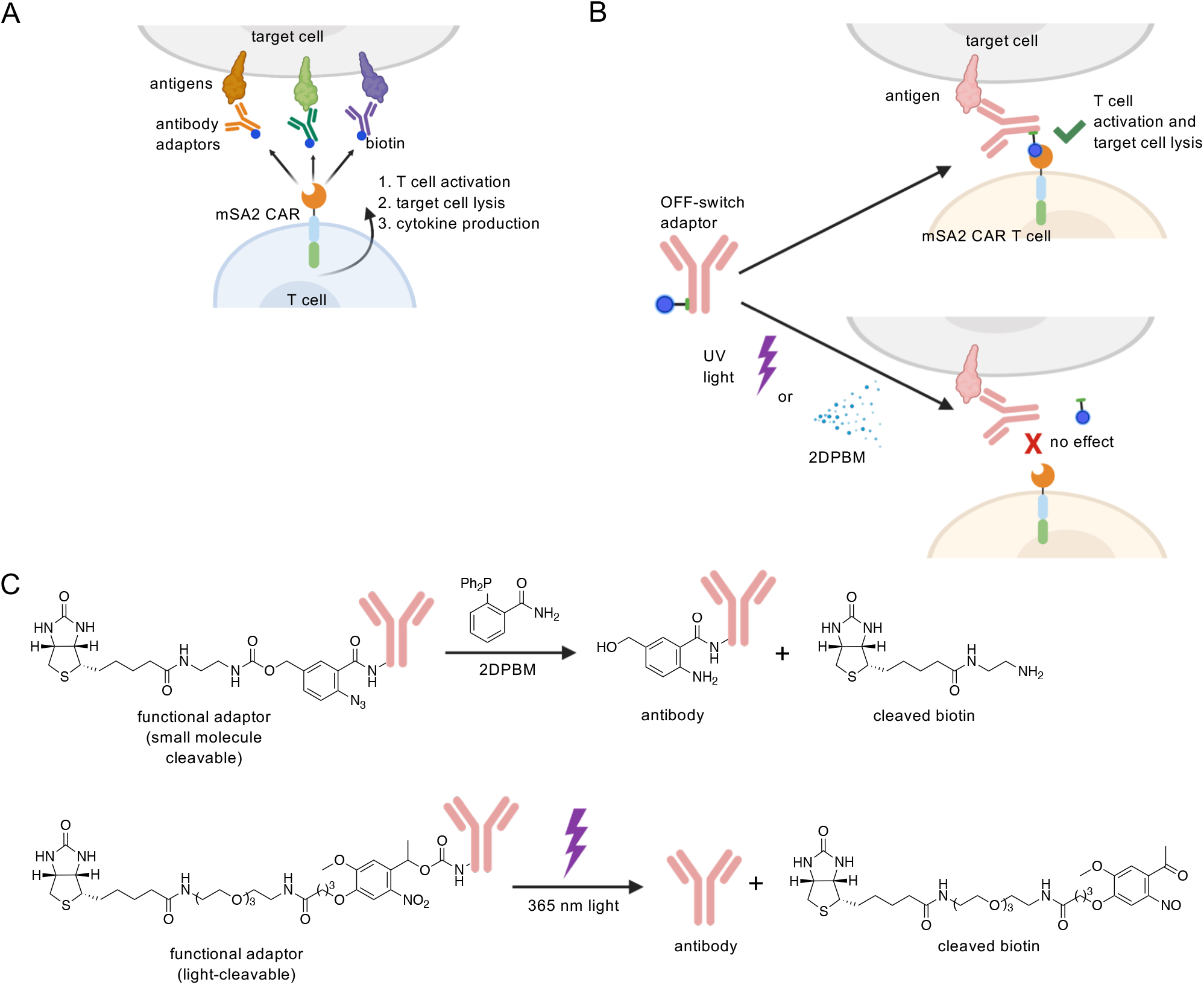
mSA2 universal CAR T cells and the OFF-switch conditional control concept. (A) mSA2 CAR T cells bind to biotinylated antibody adaptors leading to specific recognition of one or more antigens on target cells and downstream T cell activation and effector functions. (B) When antibody adaptors are exposed to UVlight or the small molecule, 2DPBM, the biotin tag is cleaved and the adaptor is incapable of mediating interaction between mSA2 CAR T cells and target cells. (C) Chemical structures of cleavable adaptor antibodies and conditional OFF-switch reaction products.

Stimulus-reactive chemical groups have been widely used to enhance the targeting and conditional control of small molecules, peptides, proteins, and nucleic acids^15^, and offer a potential means to control universal CAR T cell therapy^16^. To this end, small molecule drug systems would be advantageous as they could provide physicians with systemic control over CAR T cells through variable dosing and predictable systemic drug elimination. Small molecule OFF-switch adaptors could serve as safety switches for rapid dampening or cessation of therapy. Light is also an attractive stimulus, as it could provide precise temporal and spatial control of receptor activity using a signal that can be administered in a variety of ways, e.g., via an endoscopic fiberoptic probe^17^.

By employing stimulus-reactive chemical linkers we have generated universal OFF-switch adaptors that control CAR activity in response to UV-light and small molecule triggers. Adaptors are fitted with biotin tags attached via stimulus cleavable linkers. In the absence of the stimulus, the adaptors mediate the interaction between the CAR T cells and target cells, leading to T cell activation and target cell lysis. However, upon UV exposure inducing bond photolysis or addition of a small molecule phosphine leading to azide to amine reduction and subsequent 1,6-elimination, the biotin tags are cleaved from the adaptor, thus preventing mSA2 CAR T cells from engaging with target cells. This cleavage prevents CAR T cell functionality and therapeutic response (Figure 1B). The OFF-switch adaptor strategy presents a novel mechanism to turn off or tune universal CAR T cell activity, potentially mitigating therapeutic toxicities.

## Results

### Engineering conditional biotin OFF-switch adaptors

Seeking to gain OFF-switch control of mSA2 CAR T cells, we generated antibody adaptors conjugated to biotin via cleavable linkers (Figure 1C). To create a light-cleavable biotin, we reacted a commercially available NHS-carbonate with clinically relevant trastuzumab (anti-HER2) and rituximab (anti-CD20) antibodies and thereby created a nitrobenzyl carbamate linkage that is cleavable by exposure to 365 nm UV-light^18^. To create the small molecule-cleavable adaptors, we synthesized an NHS-ester biotin with a *para*-azido benzyl carbamate linker (Supporting Information, Figure S1), that is labile upon reduction of the azido group to an amine group through addition of the phosphine 2-(diphenylphosphanyl) benzamide (2DPBM)^19^. This NHS-ester was then reacted with rituximab and trastuzumab antibodies, stochastically modifying lysine residues at the protein surface. We also created adaptor antibodies using non-cleavable linkers consisting of two ethylene glycol moieties (PEG2). All adaptors were assessed by a 4’-hydroxyazobenzene-2-carboxylic acid (HABA) assay to quantify the number of biotin molecules per antibody, showing a range of 7-18 biotins (Supporting Information, Figure S2).

Next, we assessed the ability of UV-light to cleave the adaptor biotin on the surface of tumor cells (Figure 2A). HER2+ tumor cells were incubated with different concentrations of UV-cleavable (UVcl) or PEG2 (non-cleavable) trastuzumab adaptor or no adaptor. Unbound antibody was washed away, and cells were exposed to 365 nm light for 0-60 seconds. Cells were then washed and stained with fluorescently labeled streptavidin and analyzed by flow cytometry to quantify cell surface biotins (Figure 2B). We found that the level of cell surface biotin was rapidly reduced by UV-light exposure for the OFF-switch adaptor, decreasing by half within 15 seconds of exposure and to background staining levels (complete cleavage) within 45 seconds. Streptavidin-APC levels also correlated with the amount of adaptor used for staining. Importantly, adaptors with inert PEG2 linkers did not show reduction in streptavidin-APC, and the photochemical reduction in cell surface biotin was specific to the UV-cleavable adaptors.

**Figure 2.**
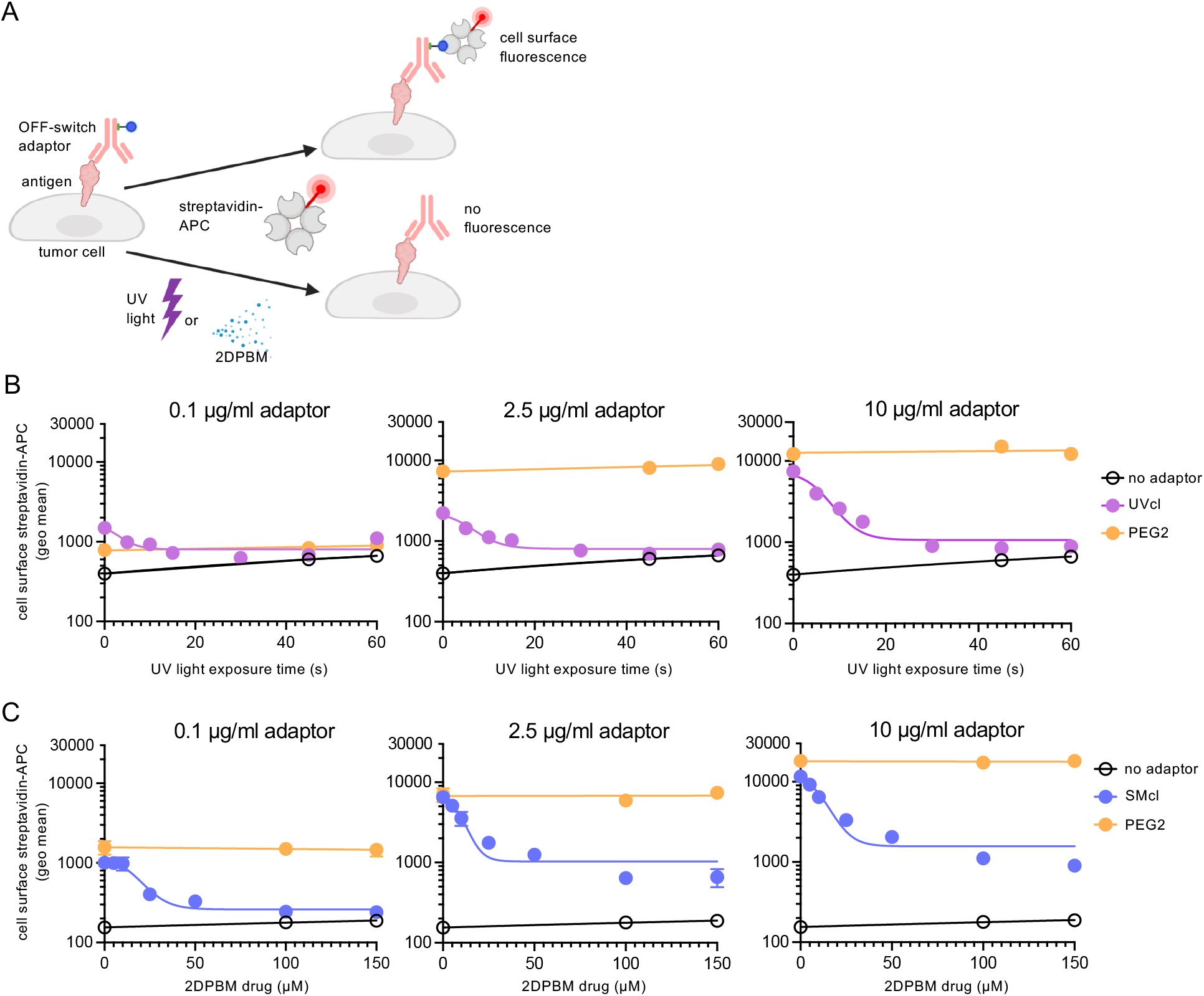
Conditional OFF-switch biotin labeling of cell surfaces by cleavable antibody adaptors. (A) Schematic representation of experimental design. Once the adaptor is bound to the target antigen on the cell surface, if cleavable adaptors are exposed UV-light or 2DPBM, the biotin tag is cleaved off thus preventing binding of streptavidin-APC to the cell surface adaptor. (B) K562-HER2 tumor cells were incubated at the indicated concentrations with UV-cleavable (UVcl) or non-cleavable PEG2 anti-HER2 adaptors and exposed to UV-light for the indicated times or (C) small molecule-cleavable (SMcl) adaptors treated with the indicated concentrations of 2DPBM, stained with streptavidin-APC and assessed by flow cytometry. Three biologically independent experiments and error bars denote ± s.d.

To test the ability of the small molecule, 2DPBM, to cleave biotin from the cleavable OFF-switch adaptors, co-incubation and analysis were carried out similarly to the above UV experiments, except that instead of UV-exposure, adaptor-labeled target cells were cultured in the presence of increasing concentrations of 2DPBM (0-100 µM) for one hour (Figure 2A). Following this incubation, cells were washed, stained, and evaluated by flow cytometry (Figure 2C). Small molecule-cleavable (SMcl) adaptors showed reduced streptavidin-APC levels with increasing concentrations of 2DPBM whereas levels with non-cleavable PEG2 adaptors were unchanged. Similar to the UVcl adaptors, APC levels correlated with the amount of adaptor applied. One difference between the two switches was that at high levels of adaptor, the 2DPBM-induced cleavage was incomplete as labeling remained significantly higher than the no adaptor control samples. However, we reasoned that the 10-fold reduction in labeling would likely be sufficient to effectively reduce mSA2 CAR T cell activity.

### UV OFF-switch adaptor control of universal mSA2 CAR T cells

Next, we tested whether the UV-cleavable adaptors were capable of mediating OFF-switch control of mSA2 CAR T cell activity. First, to generate mSA2 CAR T cells with a high level of CAR expression, we cloned the mSA2 CAR-T2A-TagBFP coding region from our previously reported lentiviral system into a gamma-retroviral vector backbone (Figure 3A)^12, 20^. When packaged into viral particles and transduced into primary human T cells, this vector yielded 50-75% expression efficiency (Supporting Information, Figure S3). HER2-positive tumor target cells were stained with varying doses of UV-cleavable adaptor, PEG2 adaptor, or no adaptor and then incubated with mSA2 CAR T cells. Co-incubations were then exposed to UV-light for different times and placed in a cell culture incubator for 24 hours. Following incubation, we analyzed CD69 and CD107a T cell activation marker expression on mSA2 CAR T cells by flow cytometry (Figure 3B). We observed that cells not exposed to UV-light, showed strong up-regulation of CD69 and CD107a markers compared to the no adaptor control. But even after only 10 seconds of UV-light exposure, CAR T cells showed significantly lower levels of both markers, indicating that UV-light adaptors could indeed lead to switching off CAR activity. Importantly, T cell activation mediated by PEG2 adaptors, and a traditional anti-CD20-CAR were unaffected by the light exposure (Supporting Information, Figure S4). We also repeated these experiments using an anti-CD20 rituximab UV-cleavable adaptor and observed conditional control of CAR T cell activation in response to CD20+ positive target cells, demonstrating that the system can be generally applicable to different adaptor antibodies and antigens (Supporting Information, Figure S5). Finally, we collected the supernatants from these co-incubation experiments and analyzed them by ELISA for production of IFNγ, a pro-inflammatory cytokine commonly up-regulated upon T cell activation. IFNγ levels mirrored that of T cell activation markers, again demonstrating OFF-switch control over mSA2 CAR T cell activation and the ability to control pro-inflammatory environmental signals mediated by the mSA2 CAR T cells (Figure 3C).

**Figure 3.**
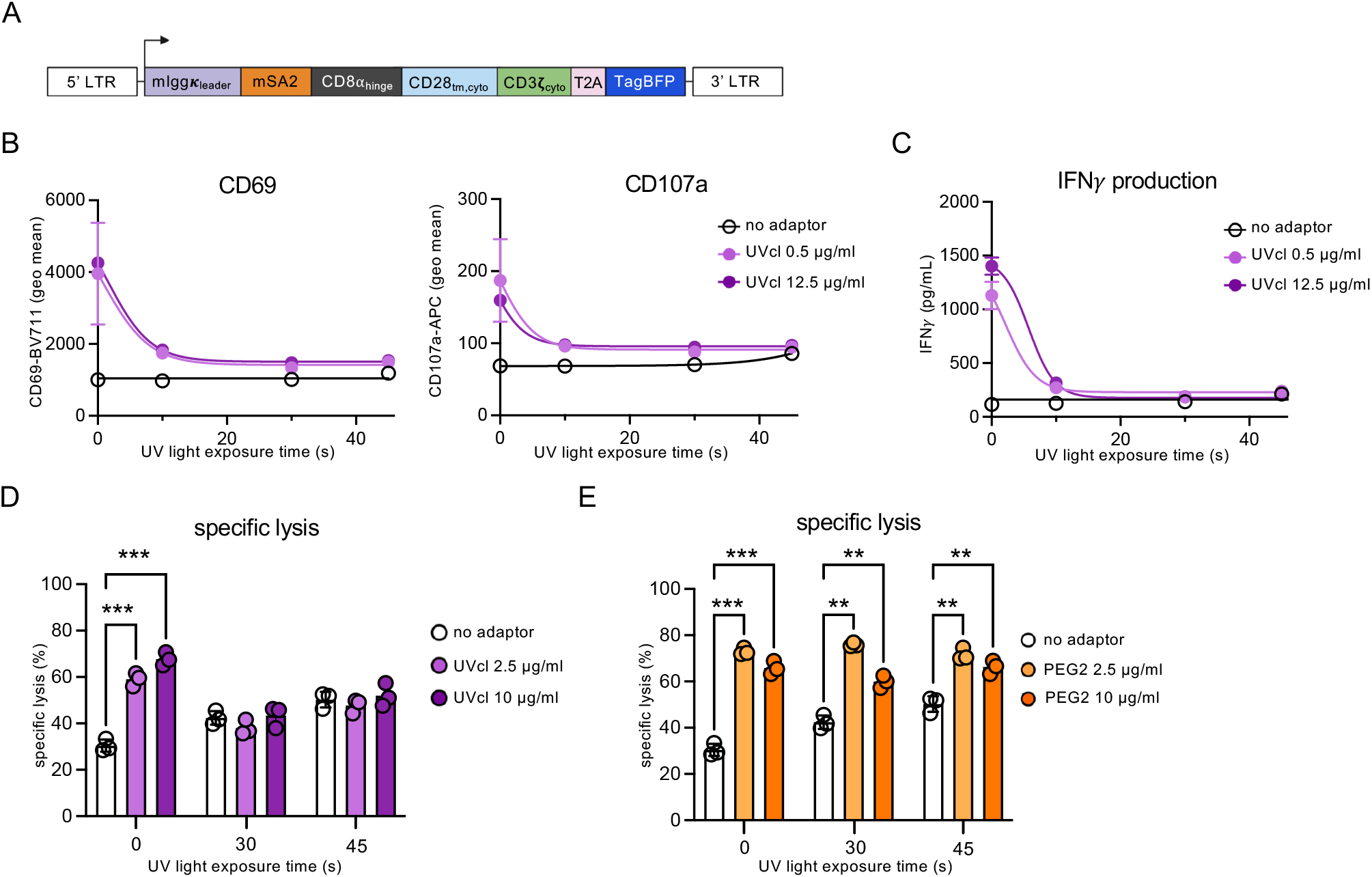
UV-cleavable adaptors mediate conditional control of mSA2 CAR T cell functions. (A) Schematic representation of the mSA2 CAR T cell construct. (B) Flow cytometry analysis of CD69 and CD107a T cell activation markers on mSA2 CAR (TagBFP+) cells co-incubated with K562-HER2 target cells. (C) ELISA for IFNψ production from primary human mSA2 CAR T effector cells co-incubated with K562-HER2 target cells and indicated concentrations of UV-cleavable adaptor. (D) Specific lysis of target cells by co-incubated primary human mSA2 CAR T cells and 2.5 or 10 µg/mL of the indicated UV-cleavable adaptors. (E) mSA2 CAR T cells were combined with target cells labeled with no adaptor or PEG2-trastuzumab at the indicated doses and exposed to UV-light for the indicated durations, incubated for 24 hours and analyzed by flow cytometry for specific lysis of the target cells. A 2-way ANOVA test was performed using Tukey’s post-hoc analysis for multiple comparisons. “**” denotes a significance of p < 0.01”***” denotes a significance of p < 0.001, three biologically independent experiments ± s.d..

UV OFF-switch adaptors were also evaluated for mediating mSA2 CAR T cell lysis of target cells in co-culture experiments by flow cytometry (Figure 3D). Significant levels of specific lysis were observed for adaptors compared to the no adaptor control, and UV-cleavable adaptor co-incubations that were exposed to UV-light, displayed a decrease in cell lysis, to levels equivalent to those of the no adaptor control. Lysis by PEG2 adaptors were unaffected by UV-light exposure (Figure 3E). Of note, with increasing UV-light exposure we observed increasing lysis of tumor cells by mSA2 CAR T cells in the absence of antibody adaptor. We concluded that this increase in target cell lysis was mediated by T cells, as the UV-light on its own did not affect the viability of the target cells (Supporting Information, Figure S6). As the light does not induce T cell activation markers, it is likely the UV-light is damaging target cells and triggering antigen independent CAR T cell lytic abilities. This result is consistent with previous reports ^21^.

Taken together, these experiments show that the UVcl adaptors can trigger CAR T cell signaling and cancer cell lysis which can be rapidly turned off by UV-light exposure. Excellent ON to OFF switching was observed by staining for cell surface biotin and could be finely tuned through titrating exposure time. The cell surface biotin levels correlated well with T cell activation levels and tumor cell lysis in cell co-incubation experiments. The approach was demonstrated for multiple antibodies and target cells, suggesting general applicability.

### Small molecule OFF-switch adaptor control of universal mSA2 CAR T cells

In order to test if the SMcl adaptors could be switched off by the small molecule 2DPBM, we stained HER2 positive tumor cells with varying doses of SMcl adaptor, PEG2 adaptor or no adaptor, and then treated these cells with 2DPBM at increasing concentrations. After a 1 hour incubation at 37 °C, mSA2 CAR T cells were added to the stained targets, and cells were co-incubated for 24 hours. The next day we analyzed CD69 and CD107a T cell activation marker expression on mSA2 CAR T cells by flow cytometry (Figure 4A). In the absence of 2DPBM, markers were strongly up-regulated in comparison to the no adaptor (negative) control and matched the non-cleavable PEG2 (positive) control. Gratifyingly, this activation was down-regulated in response to small molecule treatment. The response to 2DPBM was dose-dependent and background levels of activation were achieved at the maximum dose of 100 µM 2DPBM. Moreover, this OFF-switch activity was specific to SMcl adaptors as neither PEG2 adaptors nor a traditional anti-CD20-CAR were affected by 2DPBM (Supporting Information, Figure S7). We also repeated the same experiments by using anti-CD20 rituximab adaptors, demonstrating that the small molecule OFF-switch strategy can be applied to targeting different antigens (Supporting Information, Figure S8). Analyzing the co-incubation supernatants for IFNγ, we observed a reduction in IFNγ that correlated with 2DPBM levels, thus further demonstrating the ability of SMcl adaptors to control mSA2 CAR T cell-induced pro-inflammatory environmental signals (Figure 4B).

**Figure 4.**
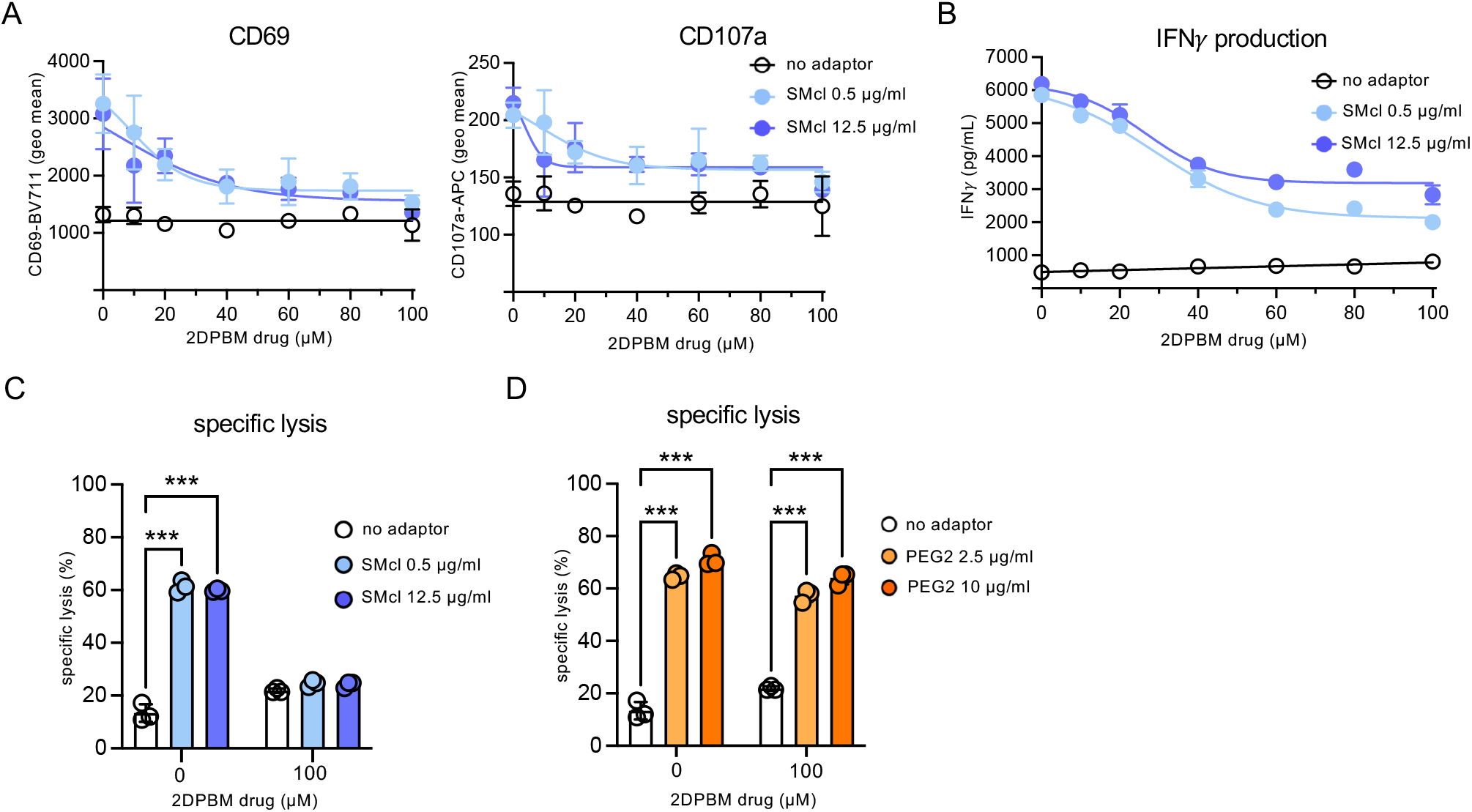
Small molecule-cleavable adaptors mediate conditional control of mSA2 CAR T cell functions. (A) Flow cytometry analysis of CD69 and CD107a T cell activation markers on the mSA2 CAR (TagBFP+) population from the co-incubations with K562-HER2 target cells. (B) ELISA for IFNγ production from primary human mSA2 CAR T effector cells co-incubated with K562-HER2 target cells and indicated concentrations of small molecule-cleavable adaptor. (C) Specific lysis of target cells by co-incubation of primary human mSA2 CAR T cells and 0.5 or 12.5 µg/mL of the indicated small molecule-cleavable adaptors. (D) mSA2 CAR T cells were combined with target cells labeled with no adaptor or PEG2-trastuzumab at the indicated doses with or without 2DPBM (100 µM), incubated for 24 hours and analyzed by flow cytometry for specific lysis of the target cells. A 2-way ANOVA test was performed “***” denotes a significance of p < 0.0001, three biologically independent experiments ± s.d.

Next, mSA2 CAR T cells and small molecule-cleavable adaptors were evaluated for target cell lysis. Cell lysis was increased when the mSA2 CAR T cells and targets were incubated with adaptors compared to the no adaptor control, and when incubated with 2DPBM, cell lysis was reduced to levels equivalent to the no adaptor control (Figure 4C). PEG2 adaptor-mediated lysis was unaffected by 2DPBM addition (Figure 4D).

Altogether, these results demonstrate the robust OFF-switch activity and specificity of the SMcl adaptors to control universal CAR T cell activity with a small molecule. The T cell activation and target cell lysis results match the staining experiments and the results obtained for the light-triggered OFF-switch. Good switching ratios were observed and the tunability of T cell function through increasing small molecule concentration showed greater sensitivity than the optical OFF-switch.

### OFF-switch adaptors mediate conditional control of NFAT-inducible transgene expression

To further investigate OFF-switch adaptor functionality, we tested whether adaptors could conditionally control transgene expression via a nuclear factor of activated T cells (NFAT) response construct. This construct consists of an NFAT-inducible promoter and the IL-2 core promoter upstream of a GFP reporter^22^. Upon T cell activation, NFAT translocates to the nucleus where it can bind to the NFAT promoter response elements (NFAT-RE) and activate GFP transcription (Figure 5A). In addition to demonstrating T cell activation, similar NFAT-inducible constructs have been used to specifically deliver therapeutic payloads, such as the IL-12 cytokine, locally to tumors^23^. We co-transduced primary human T cells with retroviruses containing the mSA2-CAR and the pNFAT-GFP construct. We then co-incubated mSA2 CAR pNFAT-GFP cells with K562-HER2 cells pre-labeled with antibody adaptors and exposed the cell mixtures to UV-light or 2DPBM. Following a 24 hour co-incubation, we assayed mSA2 CAR T cells for GFP expression by flow cytometry. mSA2 CAR T cells co-incubated with UVcl and SMcl adaptors displayed high levels of GFP induction compared to no adaptor controls, and this induction was inhibited to near background levels after the addition of either conditional stimulus (Figure 5B & C). The PEG2 adaptor showed a similarly high level of GFP induction that was unaffected by UV-light. There was an 18% decease of GFP-positive cells after 2DPBM addition, however, this level was much lower than the 42% reduction observed with the SMcl adaptor (Figure 5B & C).

**Figure 5.**
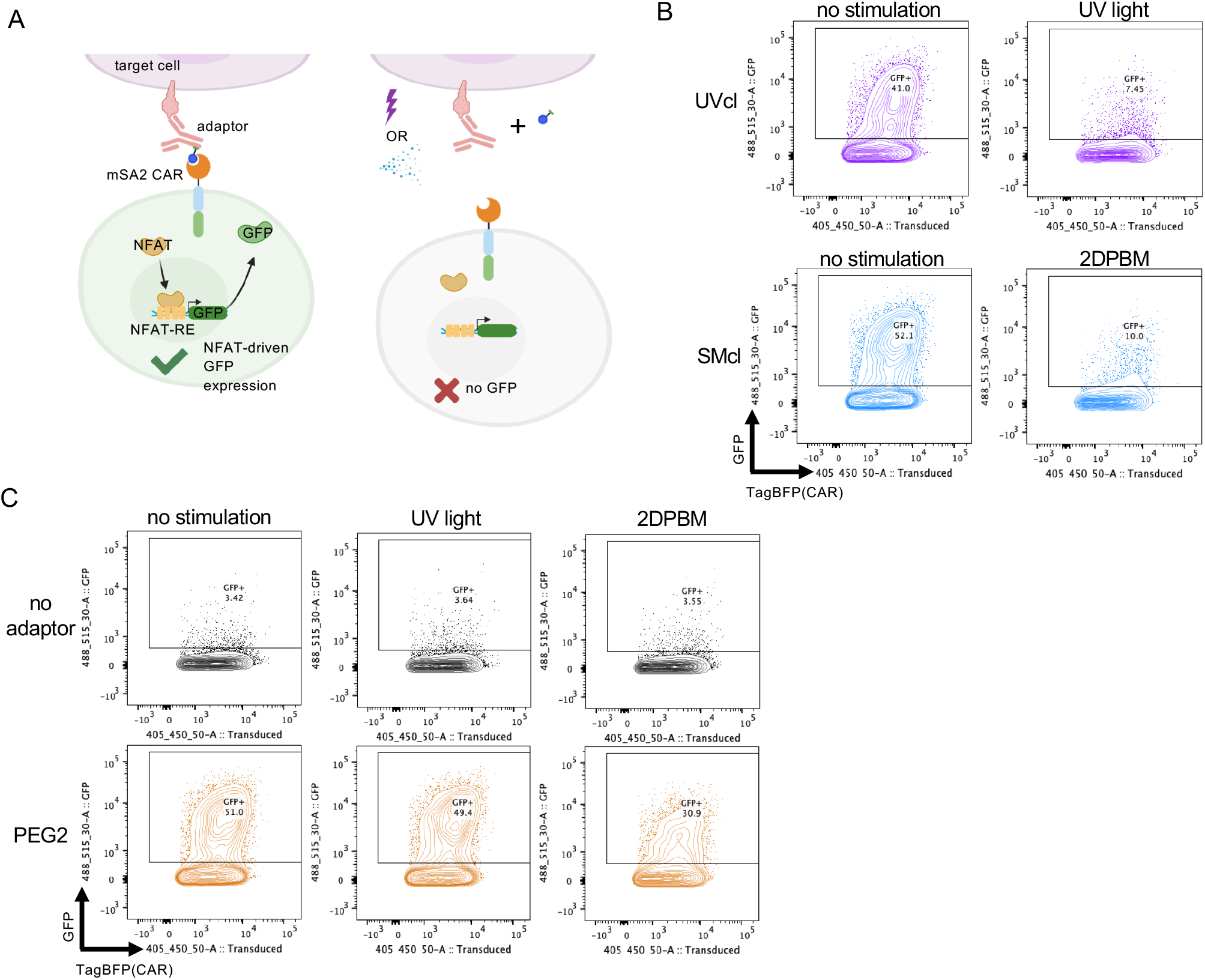
OFF-switch adaptors mediate conditional control of NFAT-inducible transgene expression. (A) Diagram of OFF-switch adaptor control of NFAT-inducible GFP. mSA2 CAR T cells bind to the adaptor on target cells leading to T cell activation and NFAT translocation to the nucleus and binding to the NFAT response elements (NFAT-RE), thus activating GFP expression. GFP expression can be prevented by adaptor cleavage. (B) Flow cytometry plots of GFP expression vs. CAR (TagBFP marker) using conditional adaptors or (C) an inert PEG2 adaptor and no adaptor controls. For (B) and (C) one representative flow cytometry plot from three replicates is shown.

### Precision multi-antigen OFF-switch control of mSA2 CAR T cells

Universal CARs uniquely enable the simultaneous targeting of multiple antigens with the same cell product, simply by adding multiple adaptors. Thus, we decided to investigate the ability of cleavable adaptor combinations to simultaneously perform multi-antigen conditional targeting. For instance, it could be desirable to target one antigen constitutively (a validated target), and to simultaneously target a second antigen (potentially a new antigen with unknown toxicities) with a conditional OFF-switch adaptor to mitigate any safety risks. To test the potential for dual targeting, we set-up co-incubations of cells positive for two different antigens together with mSA2 CAR T cells and different adaptors. To distinguish the two target cell lines (K562-HER2 and K562-CD20) we labeled them with different fluorescent dyes. We then incubated these cells with different light-or 2DBPM-treated adaptors, followed by removal of excess adaptor and addition of mSA2 CAR T cells (Figure 6A). After 24 hours we analyzed cell co-incubations for specific lysis of both K562-CD20 and K562-HER2 target cell populations by flow cytometry. This assay strategy was validated by testing individual adaptors for specific activity against antigen+ targets vs. antigen(-) ones (Supporting Information, Figure S9). The first adaptor combination that we tested was a UV-cleavable trastuzumab adaptor paired with a non-cleavable rituximab-PEG2 adaptor which would be expected to lyse target cells according to the logic operation of (HER2 AND NOT(UV)) OR (CD20) (Figure 6B). We observed that in the absence of exposing adaptors to UV-light both K562-HER2 and K562-CD20 cell lines were efficiently lysed by mSA2 CAR T cells. However, with light exposure, the K562-CD20 cells were again efficiently lysed, but K562-HER2 cells in the same well showed markedly increased survival, demonstrating UV-mediated protection specifically of this cell population. We then performed an experiment testing a second combination of adaptors, trastuzumab-SMcl and rituximab-PEG2 adaptors which would be expected to lyse target cells according to (HER2 AND NOT(2DPBM)) OR (CD20) (Figure 6C). With this combination, in the absence of adaptor exposure to 2DPBM we observed efficient cell lysis of both HER2 and CD20 positive target cell lines, while in the presence of the small molecule 2DPBM we saw the anticipated high level of lysis of K562-CD20 cells with specific down-regulation of K562-HER2 cell lysis. Finally, we tested the combination of two conditional adaptors to provide a potential means for orthogonal conditional regulation of the simultaneous targeting of two different antigens. Testing the combination of trastuzumab-SMcl and rituximab-UVcl adaptors (HER2 AND NOT(2DPBM)) OR (CD20 AND NOT(UV)), we observed cell lysis was reduced in CD20 positive target cells when the adaptor was UV-exposed, and in HER2 positive target cells when the adaptor was treated with 2DPBM with significantly higher levels of killing by each adaptor in the absence of the stimulus (Figure 6D).

**Figure 6.**
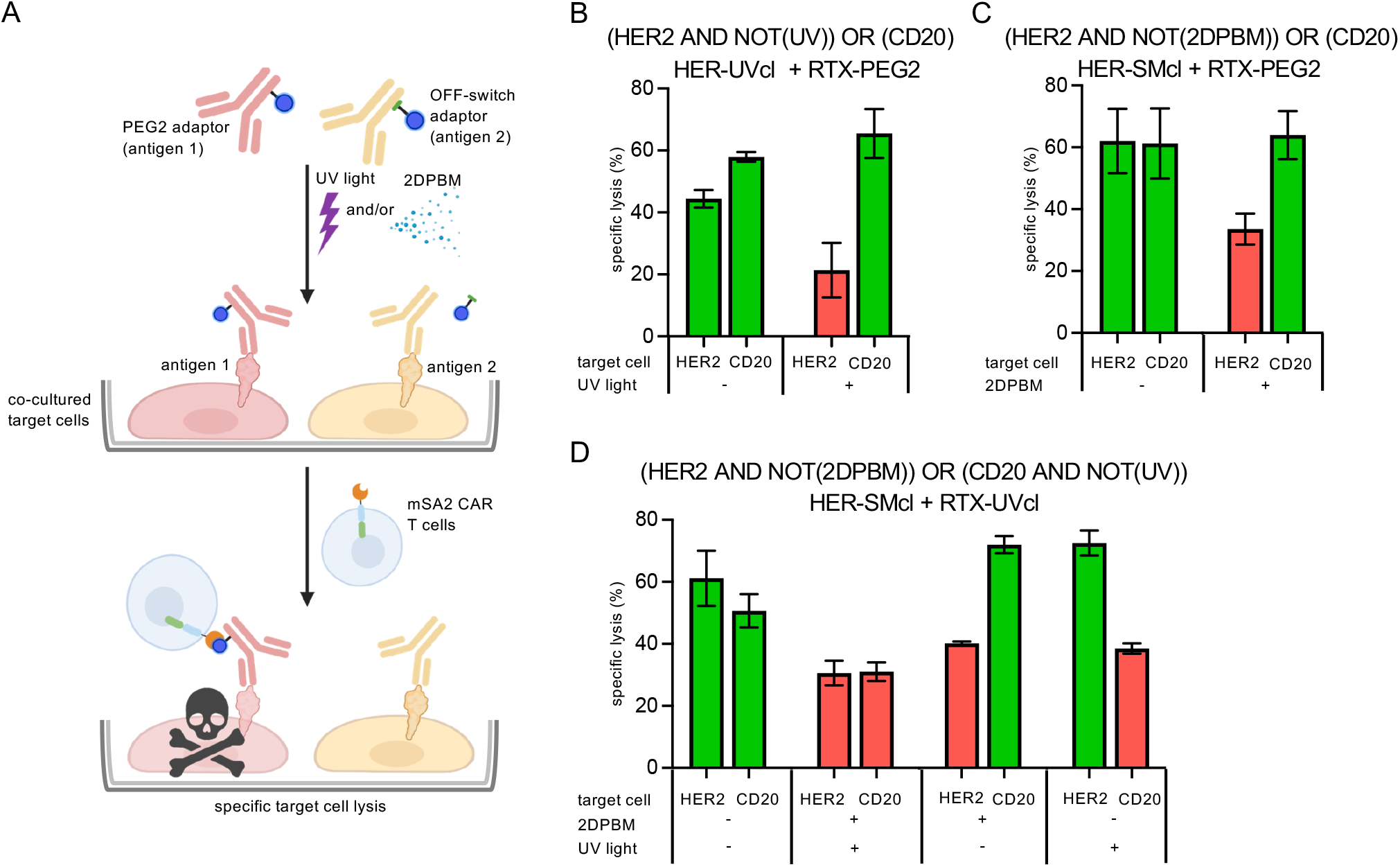
Logical programming of mSA2 CAR T cell-mediated lysis by OFF-switch adaptor combinations. (A) Schematic representation of a representative combinatorial adaptor experiment and OFF-switch behavior. (B) Specific lysis of target cell lines labeled with indicated antibody adaptors treated +/– UV-light exposure by co-incubated primary human mSA2 CAR T cells. (C) Specific lysis of target cell lines labeled with indicated antibody adaptors treated +/– 2DPBM exposure by co-incubated primary human mSA2 CAR T cells. (D) Specific lysis of target cell lines by co-incubated primary human mSA2 CAR T cells and indicated antibod adaptors that were exposed +/– UV-light and/or the 2DPBM small molecule. Green bars were used to represent conditions where lysis is expected to occur, and red bars indicate predicted conditions where lysis is expected to be turned off. n = 3 biologically-independent experiments ± s.d.

## Discussion

Cleavable OFF-switch adaptors show the ability to rapidly intercept and sensitively tune mSA2 CAR T cell activity. For the UV-cleavable adaptor, experiments measuring biotin on the surface of target cells, showed that biotin was rapidly removed, with a half-life of 10 seconds of UV exposure and a near complete cleavage at 30 seconds. This rapid control of cell surface biotin translated to control of CAR T cell activation and target cell lysis. Fast-acting prevention of CAR activity using the UV-cleavable OFF-switch adaptors could potentially be used to protect sensitive anatomical regions from universal CAR T cells while allowing therapy to proceed in other areas, thus preventing ON-target/OFF-tumor toxicities. The UV-cleavable adaptors could be interfaced with existing methods to administer light in a surgical setting using existing methods to provide precise spatiotemporal control of receptor activity^17, 24^. While the system showed robust OFF-switch functionality, we observed enhanced antigen-independent toxicity of the CAR T cells in response to UV-light. This result suggests that one area for technical advance would be in designing light cleavable adaptors responsive to longer wavelengths of light that are potentially less likely to lead to cellular damage^25^. The complementary small molecule adaptors also displayed sensitive OFF-switch control of cell surface biotin and mSA2 CAR T cell activity. SMcl adaptors showed tunable deactivation in response to 2DPBM, reaching an IC_50_ at 25 µM and near total inhibition at 100 µM. Receptor tuning is essential for addressing ON-target/OFF-disease toxicity from overactive signaling, which currently leads to many patients (73-85%) experiencing serious toxicities including cytokine storm, neurotoxicity, or CAR T related encephalopathy syndrome^7^. While traditional antibody adaptors allow for some level of tuning with adaptor dose, the circulating half-life of antibody drugs can be up to multiple months, and thus the small molecule cleaving system could provide an advantageous method to systemically prevent further toxic activity from mSA2 CAR T cells without negatively impacting the CAR T cells. This method would compare well to cell ablative approaches such as drug-controlled cell suicide switches, as it would allow for restarting CAR T cell activity through future administration of the same of a different adaptor^26^.

Adaptor combination experiments demonstrated the unique ability of cleavable adaptors to mediate simultaneous multi-antigen OFF-switch control. Combining either a trastuzumab-UVcl or SMcl adaptor with an inert rituximab adaptor allowed for selective deactivation of antigen targeting of the first antigen (HER2) while permitting continued targeting of the second (CD20). This enhanced programmability could allow for the cessation of targeting an experimental, problematic antigen while continuing to allow therapeutic targeting of a safer, established antigen. Furthermore, combining a rituximab-UVcl adaptor with a small molecule cleavable adaptor targeting HER2 (trastuzumab-SMcl) allowed for dual stimulus OFF-switch control over CAR activity. The versatility and precision control of the OFF-switch adaptor technology compares well to existing drug-switchable receptor systems, including for universal CAR T cells, where the creation of multiple receptors would be required^27^.

In addition to providing a robust means of OFF-switch control of mSA2 CAR T cells with potential clinical utility, these results more broadly validate the mechanism of reactive chemical linkers to control universal CAR activity. A similar approach could be generalized to other universal CARs that bind to adaptors with alternative chemical tags (such as fluoresceine, benzylguanine, PNE, etc.)^12, 13^. Additionally, the cleavable OFF-switch concept could be applied to creating adaptors responsive to other stimuli including, alternative wavelengths of light, other small molecules, and endogenous cellular signals (e.g., enzymatic function)^28^. Cleavable adaptors provide a novel approach for accomplishing dynamic and precise OFF-switch control of antigen targeting by cellular therapeutics

## Methods

*Production of antibody adaptor conjugates*. Rituximab (Rituxan, Genentech) and trastuzumab (Herceptin, Genentech) were first buffer exchanged into PBS using 5 mL 7K MWCO Zeba Spin Desalting Columns (ThermoFisher Scientific). Antibodies were then co-incubated with a 20-fold molar excess of NHS ester: PC-biotin-NHS ester (Click Chemistry Tools), Biotin-PEG2-NHS ester (BroadPharma), or SMcl-NHS ester for 30 minutes at room temperature, followed by buffer exchange into PBS using 5 mL 7K MWCO Zeba Spin Desalting Columns.

### mSA2 CAR expression vector and gamma retrovirus production

The MSGV1 and RD114 retroviral plasmids were a gift of Dr. U. Kammula. The mSA2 CAR was cloned from the previously generated pHR construct into the MSGV1 backbone via isothermal assembly into the NcoI and NotI restriction enzyme sites. pSIRV-NFAT-eGFP was a gift from Peter Steinberger (Addgene plasmid #118031; http://n2t.net/addgene:118031; RRID:Addgene_118031). Retrovirus was produced following established methods^12, 29^.

### Cell line culture

Human tumor cell lines K562+CD20 and K562+HER2 stably expressing CD20 and HER2, respectively, were generated by transducing K562 (ATCC, CCL-243) or K562+ZsGreen-FF-luciferase cells with lentiviral particles encoding human CD20 (UniProt: P11836) or human HER2 (UniProt: P04626) coding regions. One week after transduction cells were stained for HER2 or CD20 with fluorescently labeled antibodies and underwent FACS for high antigen expression. Cells were periodically assayed for stable antigen expression by flow cytometry. HEK293T cells (ATTC, CRL-3216), used for lentivirus production were cultured at 37°C in DMEM supplemented with 10% FBS, and Penicillin-Streptomycin.

### Primary human T cell culture and retroviral transduction

mSA2 CAR T cells were generated by first isolating PBMCs from deidentified human Buffy Coat samples purchased from the Pittsburgh Central Blood Bank fulfilling the basic exempt criteria 45 CFR 46.101(b)(4) in accordance with the University of Pittsburgh IRB guidelines. PBMCs from healthy donors, were isolated using Ficoll gradient centrifugation and human T cells were obtained by magnetic sorting using the eHuman Pan T cell isolation kit (Miltenyi Biotec). Once isolated, T cells were cultured in RPMI (Gibco) supplemented with 10% Human AB serum (Gemini Bio Products) with the addition of 100 U/ml human IL-2 IS (Miltenyi Biotec), 1 ng/ml IL-15 (Miltenyi Biotec), and 4mM L-Arginine (Sigma Aldrich). HCl was added to bring the mdia to pH 7.2. T cells were activated with TransAct Human T cell activation reagent (Miltenyi Biotec) and 100 U/ml human IL-2 IS (Miltenyi Biotec), 1 ng/ml IL-15 (Miltenyi Biotec). After 48 hours, T cells were transduced with retrovirus. Briefly, polystyrene plates were coated overnight with 20 µg/mL retronectin (Takara Bio.) in PBS at 4°C. On the day of transduction, retronectin solution was removed and 2 ml of viral supernatant was added to each well and spun at 2000xg for 2 hours at 32°C. After removing 2 ml of supernatant, 1×10^6^ cells in 4ml of media were added per well and spun at 32°C for 10 mins at 1000xg and returned to the incubator. Cells were maintained every 2-3 days at 1M/cells/mL in media supplemented with IL-2 and IL-15.

### Flow cytometry staining

Cells were washed and resuspended in flow cytometry buffer (PBS + 2% FBS) and then stained using the indicated antibodies and concentrations for 30 minutes at 4°C followed by two washes with flow cytometry buffer. Live cells and singlets were gated based on scatter. mSA2 CAR+ T cells were gated based on TagBFP reporter gene expression.

### mSA2 CAR T cell activation assay

The indicated target cells were stained with Cell Trace Yellow following the manufacturer’s recommended protocol (ThermoFisher). Cells were then stained with the indicated antibody adaptors/amounts and were washed twice with flow cytometry buffer and resuspended in cell culture media. 10,000 target cells per well were then co-cultured with 50,000 mSA2 CAR T cells (effector:target=5:1) in a 96 well V-bottom plate. All co-incubation experiments with mSA2 CAR T cells were performed in DMEM media supplemented with 10% FBS. For UV-cleaving experiments, wells were then exposed to 365 nm light using a 365 nm LED (Mouser, 416-LST101G01UV01). For small molecule cleaving experiments, the indicated amounts of 2DPBM were added to wells containing all cell populations. Cells were then incubated at 37°C for 24 hours. Cells were then evaluated stained with fluorescently labeled antibodies recognizing T cell activation markers: CD69-BV711 (BD Biosciences) and CD107a-APC (BD Biosciences) and GFP for NFAT-induction experiments. Marker expression was specifically evaluated on mSA2 CAR+ by gating for the TagBFP+ population. Supernatants from primary cell assays were also collected and analyzed for IFNɣ by ELISA (BioLegend).

### Target cell lysis assay

Cell co-incubations were prepared as above for cell activation experiments except following 24 hour incubation, cells were stained with Ghost Dye Red Viability Dye (Tonbo Biosciences) to calculate dead cells, and co-incubations were analyzed by flow cytometry. Target cells were identified by Cell Trace Yellow or Cell Tracker and lysed target cells were identified by positive Ghost Dye staining. Percent specific cytotoxicity of target cells was calculated by the equation: 100*(% experimental lysis – % target-only lysis) / (100 – % target-only lysis).

### Combinatorial target cell lysis assays

For combinatorial adaptor assays, K562+CD20 and K562+HER2 target cells were stained with either Cell Trace Yellow or cell Trace Far Red using manufacturer’s protocols. Cells were then stained with antibody adaptors that were unexposed or pre-exposed to adaptor stimuli (100µM 2DPBM for 2 hours, or 2 minutes of UV-light exposure). Stained cells were then incubated with mSA2 CAR T cells for 24 hours in the cell incubator and evaluated by flow cytometry for specific lysis. The two different target cell populations were identified by Cell Trace Yellow or Cell Tracker Deep Red, and lysed target cells were identified by positive Ghost Dye staining. Percent specific cytotoxicity of each target cell population was calculated by the equation: 100*(% experimental lysis – % target-only lysis) / (100 – % target-only lysis).

### Statistical methods and data analysis

The number of replicates, mean value, and error are described in the respective figure legends and/or methods. Error bars are shown for all data points with replicates as a measure of variation within a group. Flow cytometry data was analyzed using FlowJo v10.8.1 (FlowJo, LLC), and data for all figures was presented and analyzed using Graphpad Prism v9 (GraphPad Software, LLC).

### Data availability

All data generated and analyzed during this study are included in this published article and its Supporting Information or are available from the corresponding author upon reasonable request.

## Acknowledgments

This work was supported by NIH grant R01GM142007 (J.L., A.D.). This work benefitted from using the SPECIAL BD LSRFORTESSA funded by NIH 1S10OD011925-01. This project also used the Hillman Cytometry Facility that is supported in part by award P30CA047904. Cartoons in Figures 1, 2A, 3A, 5A, and 6A were created with BioRender.com.

## Author contributions

J.L. and A.D. designed the research. M.K., E.R., Y.T., V.S., and A.B.P. carried out the experiments and analyzed the data. J.L. and A.D supervised the work. M.K., E.R., Y.T., V.S., A.B.P., J.L., andA.D. interpreted the data. J.L., E.R., M.K., and A.D. wrote the manuscript with input from all authors.

## Materials & correspondence

Correspondence and requests for materials should be addressed to J.L and A.D.

## Supporting information

Supplementary figures showing chemical synthesis of the SMcl OFF-switch adaptor, adaptor biotin quantification, mSA2 CAR expression, T cell subset analysis, functional data for rituximab adaptors, cell viability following UV exposure, and adaptor combination lysis control experiments. Supplementary methods describing chemical synthesis of the SMcl OFF-switch adaptor and supplementary references.

