## Supporting information for "Conditional control of universal CAR T cells by cleavable OFF-switch adaptors"

Electronic Supporting Information for

Jason Lohmueller, PhD

University of Pittsburgh School of Medicine

UPMC Hillman Cancer Center

5117 Centre Ave

Pittsburgh, PA 15232, USA

Alexander Deiters, PhD

University of Pittsburgh

Chevron Science Center

219 Parkman Avenue

Pittsburgh, PA 15260, USA

**Supplementary Figures**

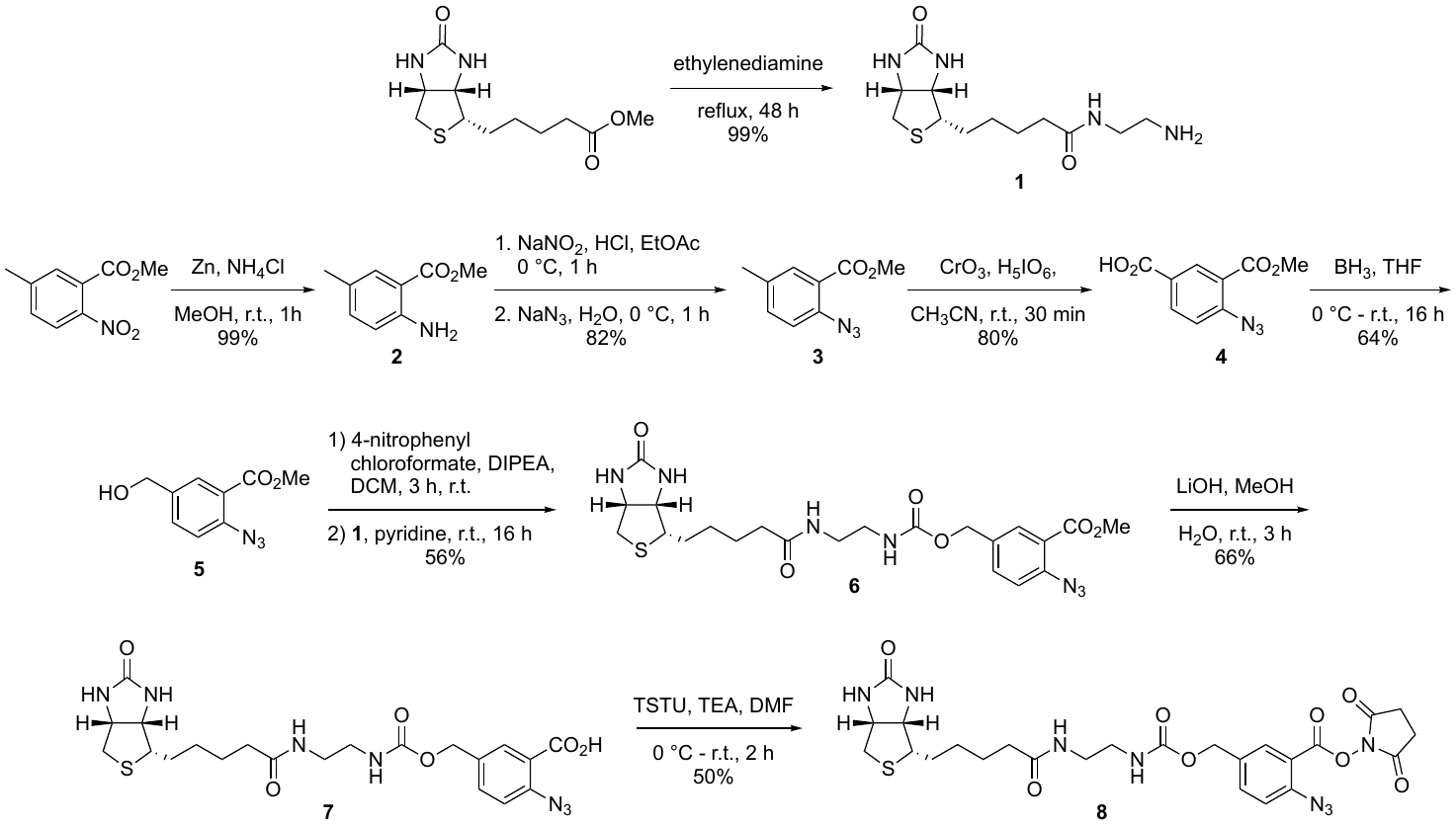

**Figure S1 | Synthesis of the SMcl OFF-switch adaptor.** Commercially available 5-methyl-2-nitrobenzoic acid methyl ester was reduced to the aniline **2** and a subsequent Sandmeyer reaction generated the aryl azide **3**. Chromate oxidation then delivered the acid **4** which was reduced to the benzylic alcohol **5**. The alcohol was coupled to the amine-modified biotin **1**, which itself was generated from biotin methyl ester, to give the phosphine cleavable ester **6**. Base-mediated methyl ester hydrolysis produced the acid **7**, which was then activated as the NHS ester **8**.

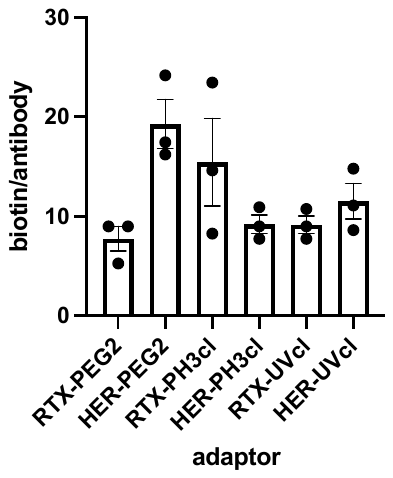

**Figure S2 | HABA assay to quantify the number of biotin molecules per antibody.** Following manufacture’s procedure, coulometric test was used to quantify the number of biotins on the adaptor. n = 3 biologically-independent experiments ± s.e.m.

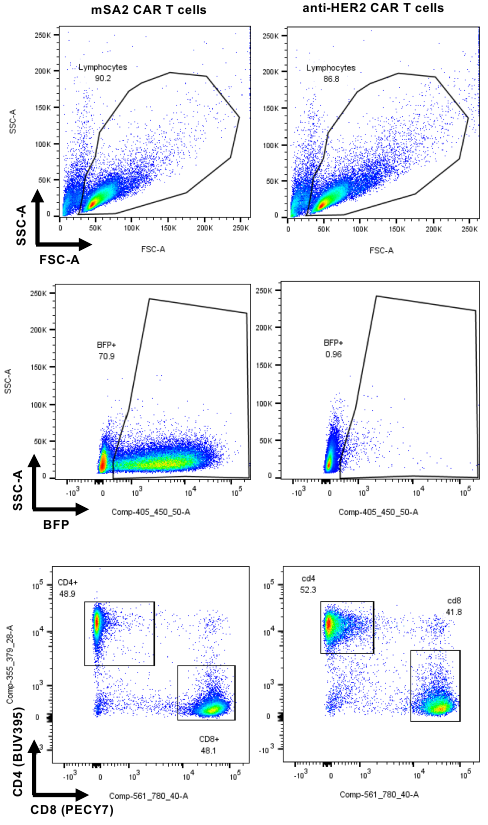

**Figure S3 | mSA2-CAR expression and characterization of T cell subsets.** At day 5 post viral transduction, primary human msa2 CAR T cells and a negative control HER2-CAR were stained with the indicated antibodies (anti-CD4-BUV395 and anti-CD8-PECY7), washed, and analyzed by flow cytometry for T cell markers and TagBFP for CAR expression.

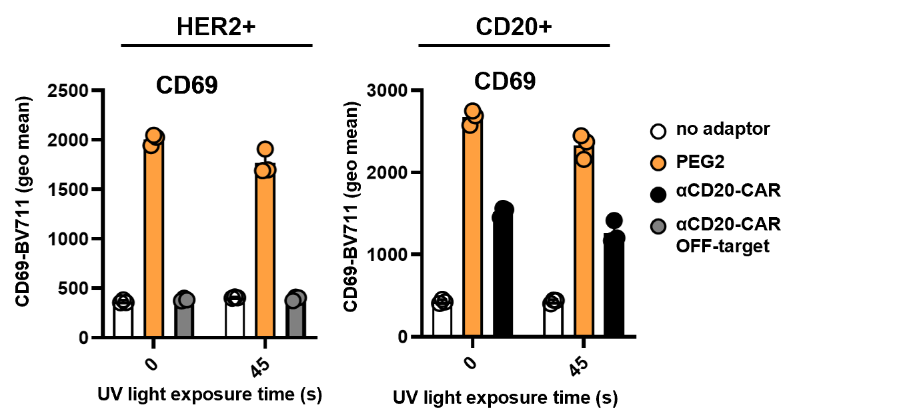

**Figure S4 | UV light exposure has no effect on CD69 T cell activation marker expression on control CAR T cells.** Flow cytometry analysis of CD69 T cell activation marker expression on the CAR T cells co-incubated with either K562-HER2 or K562-CD20 target cells and mSA2- CAR T cells with antigen specific-adaptors with inert PEG2 linker or traditional anti-CD20 CAR T cells. 2-way ANOVA test was performed using Tukey’s post-hoc analysis for multiple comparisons. “ * ” denotes a significance of p < .001, (no significant differences were observed), n = 3 biologically-independent experiments ± s.d.

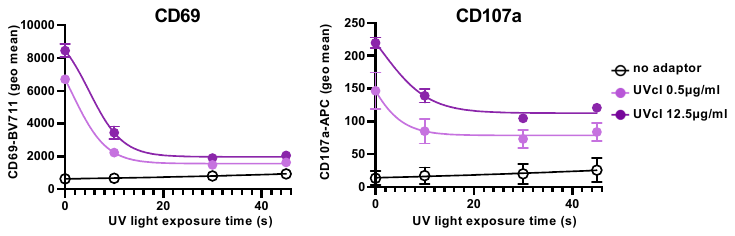

**Figure S5 | Rituximab UV-cleavable adaptor control of mSA2 CAR T cell function.** Flow cytometry analysis of CD69, and CD107a T cell activation markers on the mSA2-CAAR T cells (TagBFP+) population from 24 hours co-incubations of mSA2 CAR T cells with adaptor labeled K562-CD20 target cells exposed to UV-light for the indicated times. n = 3 biologically-independent experiments ± s.d.

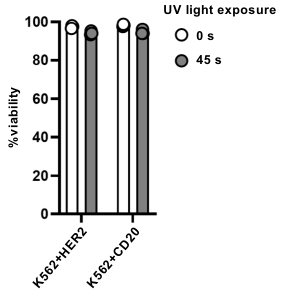

**Figure S6 | UV-light exposure has no effect on K562 target cell viability.** Indicated target cells were exposed to UV-light for 0 or 45 seconds, incubated for 24 hours and then assayed by flow cytometry for cell viability. A paired Student’s t-test was performed to determine significance. “*” denotes a significance of p < .0001 (no significant differences were observed), n = 3 biologically-independent experiments ± s.d.

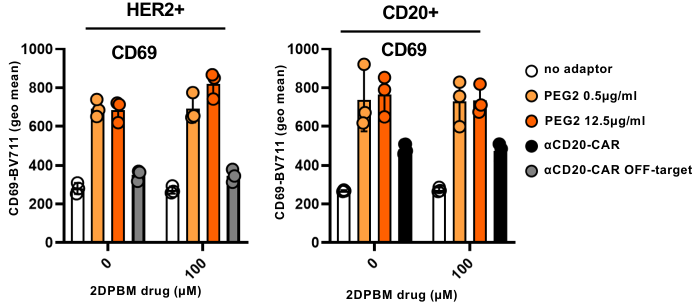

**Figure S7 | CD69 expression in mSA2 CAR T cells with PEG2-Herceptin and CD20 CAR T cells with 2DPBM exposure.** Flow cytometry analysis of CD69 T cell activation marker on the mSA2 population from the co-incubations with K562-HER2 and K562-CD20 target cells with PEG2 specific Abs and comparing to CD20 CAR T cell with different concentration of 2DPBM. 2-way ANOVA test was performed “ * ” denotes a significance of p < .0001 (no significant differences were observed), n = 3 biologically-independent experiments ± s.d.

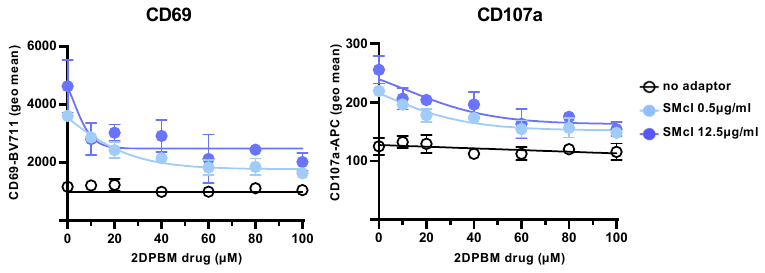

**Figure S8 | Rituximab UV-cleavable adaptor control of mSA2 CAR T cell function.** Flow cytometry analysis of CD69, and CD107a T cell activation markers on mSA2 CAR T cells (TagBFP+) from 24 hours co-incubations of mSA2 CAR T cells with adaptor labeled K562-CD20 target cells exposed to the indicated concentrations of 2DPBM. n = 3 biologically-independent experiments ± s.d.

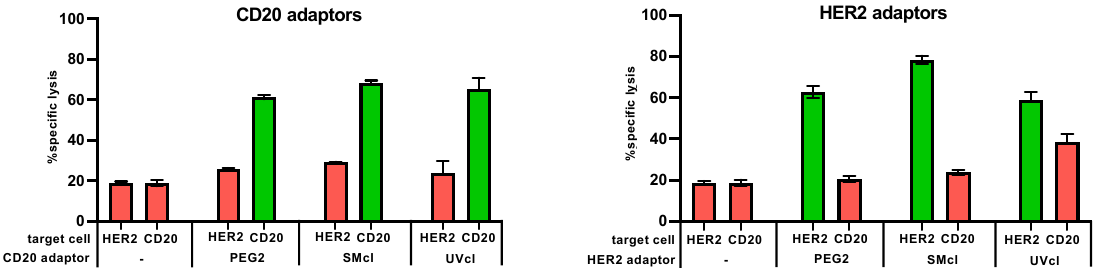

**Figure S9 | Logical programming of mSA2-CAR T cell-mediated lysis by OFF-switch adaptor combinations.** Specific lysis of the indicated target cell lines labeled with various with HER2 (trastuzumab) or RTX (rituximab) antibody adaptors that were pre-incubated +/- 2DPBM and/or +/- UV-light exposure by co-incubated primary human mSA2 CAR T cells.

**Supplementary Methods**

***N*-(2-Aminoethyl)-5-((3aS,4S,6aR)-2-oxohexahydro-1H-thieno[3,4-d]imidazol-4-yl)pentanamide (1).** The biotin methyl ester (1.50 g, 5.81 mmol) was dissolved in a MeOH (25 mL) and ethylene diamine (25 mL) solution. The reaction mixture was heated to 60 °C under reflux and stirred for 48 h. It was then concentrated under reduced pressure and the residue was suspended in toluene (3 mL), followed by evaporation of all volatiles. This was repeated a total of three times in order to remove water. No further purification was necessary and the product **47** was obtained in 99% yield (1.64 g) as a white solid. Analytical data matched those previously reported.^1^

**2-Amino-5-methyl-benzoic acid methyl ester (2).** 2-Nitro-5-methylbenzoic acid methyl ester (1 g, 5.12 mmol) was dissolved in MeOH (100 mL). Zinc dust (3.3 g, 51.2 mmol) and ammonium chloride (1.4 g, 25.6 mmol) were added and the reaction mixture was vigorously stirred at room temperature for 1 h. It was then filtered through celite (20 g) and concentrated under reduced pressure onto silica gel (2 g). The product was then purified by flash chromatography on silica gel using 20% EtOAc in hexanes as the eluent. Compound **2** was obtained in 99% yield (840 mg) as a clear solid. Note: palladium catalyzed hydrogenation would likely work as well. Analytical data matched those previously reported.^2^

**2-Azido-5-methyl-benzoic acid methyl ester (3).** Compound **2** (840 mg, 5.08 mmol) was dissolved in EtOAc (8.6 mL). The solution was cooled to 0 °C and concentrated HCl (1.6 mL) was added. In a separate vial, sodium nitrite (421 mg, 6.10 mmol) was dissolved in water (4 mL). It was then added dropwise to the solution containing compound **2** and the reaction mixture was stirred for 10 min at 0 °C. Sodium azide (397 mg, 6.10 mmol) was dissolved in water (4 mL). It was then added dropwise to the reaction mixture, which was stirred for an additional 1 h at 0 °C. The reaction mixture was slowly diluted with saturated sodium bicarbonate (30 mL). The aqueous layer was then extracted with EtOAc three times (30 mL each). The combined organic layer was dried over sodium sulfate (1 g), filtered, and concentrated under reduced pressure. No further purification was required. Compound **3** was obtained in 82% yield (800 mg) as a clear solid. Analytical data matched those previously reported.^3^

**2-Azido-5-carboxy-benzoic acid methyl ester (4).** Periodic acid (3.72 g, 16.2 mmol) was suspended in acetonitrile (40 mL) and vigorously stirred for 10 min. CrO_3_ (77 mg, 0.81 mmol) was added. Compound **3** (775 mg, 2.1 mmol) was dissolved in acetonitrile (1 mL) and added dropwise. A white precipitate immediately formed and the reaction mixture was stirred for an additional 30 min. It was then dilute with EtOAc (100 mL) and washed with 0.1 M HCl (100 mL). The layers were separated and the aqueous phase was extracted once more with EtOAc (100 mL) and the combined organic layer was washed with brine (100 mL). The organic layer was then dried over sodium sulfate (2 g), filtered, and concentrated under reduced pressure onto silica gel (1 g). The product was then purified by flash chromatography on silica gel using 1% AcOH, 20% EtOAc in hexanes followed by 1% AcOH, 65% EtOAc in hexanes as the eluent. Compound **4** was obtained in 80% yield (661 mg) as a white solid. ^1^H NMR (500 MHz, CDCl_3_) δ 12.24 (s, 1H), 8.29 (d, *J* = 7.5 Hz, 1H), 8.10 (s, 1H), 7.45 (d, *J* = 7.5 Hz, 1H), 3.90 (s, 3H). ^13^C NMR (133 MHz, CDCl_3_) δ 170.0, 165.2, 134.8, 134.6, 130.1, 129.2, 122.7, 50.1. HRMS calculated for [M-H]^-^ 220.0437, found 220.0440.

**2-Azido-5-methanol benzoic acid methyl ester (5).** Compound **4** (250 mg, 1.13 mmol) was suspended in THF (4.5 mL) and cooled to 0 °C. BH_3_ (1 M in THF, 4.5 mL, 4.5 mmol) was added dropwise and the reaction mixture was stirred for 16 h in a sealed vial, and was allowed to slowly warm to room temperature. The reaction mixture was again cooled to 0 °C and slowly diluted with saturated sodium bicarbonate (40 mL). The aqueous layer was extracted three times with DCM (40 mL each) and the combined organic layer was dried over sodium sulfate (1 g), filtered, and concentrated under reduced pressure onto silica gel (500 mg). The product was then purified by flash chromatography on silica gel using 40% EtOAc in hexanes as the eluent. Compound **5** was obtained in 64% yield (150 mg) as a clear oil. Note: some of the methyl ester was reduced as well. A better yield may be achieved by maintaining 0 °C and shortening the reaction time. ^1^H NMR (500 MHz, CDCl_3_) δ 7.88 (s, 1H), 7.64 (d, *J* = 8 Hz, 1H), 7.43 (d, *J* = 8 Hz, 1H), 4.55 (s, 2H), 3.88 (s, 3H). ^13^C NMR (133 MHz, CDCl_3_) δ 165.7, 143.8, 130.2, 129.4, 129.3, 129.1, 122.8, 64.5, 50.3. HRMS calculated for [M-H]^-^ 206.0644, found 206.0648.

**Methyl 2-azido-5-((((2-(5-((3aS,4S,6aR)-2-oxohexahydro-1H-thieno[3,4-d]imidazol-4-yl)pentanamido)ethyl)carbamoyl)oxy)methyl)benzoate (6).** Compound **5** (80 mg, 0.38 mmol) was dissolved in DCM (2 mL). DIPEA (200 µL, 1.15 mmol) was added, followed by 4-nitrophenylchloroformate (115 mg, 0.57 mmol), and the reaction mixture was stirred for 3 h at room temperature. The biotin amine **1** (218 mg, 0.76 mmol) was dissolved in pyridine (2 mL). It was then added to the reaction mixture, which was stirred for an additional 16 h at room temperature. The reaction mixture was concentrated under reduced pressure onto silica gel (200 mg). The product was then purified by flash chromatography on silica gel using 10% MeOH in DCM as the eluent. Compound **6** was obtained in 56% yield (110 mg) as a white solid. ^1^H NMR (500 MHz, CD_3_OD) δ 8.01 (s, 1H), 7.70 (d, *J* = 7.5 Hz, 2H), 7.54 (d, *J* = 7.5 Hz, 2H), 5.11 (s, 2H), 4.45-4.61 (m, 2H), 4.02 (s ,3H), 3.36-3.74 (m, 5 H), 2.99-3.23 (m, 2H), 2.24 (t, *J* = 7 Hz, 2H), 1.61-1.69 (m, 6H). (133 MHz, CDCl_3_) δ 179.1, 166.8, 166.3, 157.9, 140.4, 130.8, 129.9, 129.7, 123.2, 68.8, 64.1, 63.1, 58.9, 55.8, 40.4, 40.3, 39.0, 38.0, 28.1, 22.5, 22.3. HRMS calculated for [M-H]^-^ 518.1900, found 518.1903.

**2-Azido-5-((((2-(5-((3aS,4S,6aR)-2-oxohexahydro-1H-thieno[3,4-d]imidazol-4-yl)pentanamido)ethyl)carbamoyl)oxy)methyl)benzoic acid (7).** Compound **6** (60 mg, 0.12 mmol) was dissolved in MeOH (4.8 mL). LiOH (2 M in water, 1.2 mL, 2.4 mmol) was added and the reaction mixture was stirred for 3 h at room temperature. The reaction was quenched with AcOH (6 drops). The reaction mixture was concentrated under reduced pressure. The residue was suspended in toluene (1 mL) and concentrated under reduced pressure a total of three times. The product was then purified by column chromatography on silica gel using 1% AcOH and 12% MeOH in DCM. Compound **7** was obtained in 66% yield (40 mg) as a white solid. ^1^H NMR (500 MHz, CD_3_OD) δ 7.95 (s, 1H), 7.64 (d, *J* = 7.5 Hz, 2H), 7.44 (d, *J* = 7.5 Hz, 2H), 5.02 (s, 2H), 4.32-4.51 (m, 2H), 3.31-3.72 (m, 5 H), 2.91-3.14 (m, 2H), 2.09 (t, *J* = 7 Hz, 2H), 1.59-1.66 (m, 6H). (133 MHz, CDCl_3_) δ 179.0, 166.4, 166.3, 157.6, 140.1, 130.2, 129.4, 129.0, 122.5, 68.3, 63.6, 62.7, 58.0, 39.4, 39.2, 38.8, 37.5, 27.6, 22.0 21.6. HRMS calculated for [M-H]^-^ 504.1744, found 504.1741.

**2,5-Dioxopyrrolidin-1-yl 2-azido-5-((((2-(5-((3aS,4S,6aR)-2-oxohexahydro-1H-thieno[3,4-d]imidazol-4-yl)pentanamido)ethyl)carbamoyl)oxy)methyl)benzoate (8).** Compound **7** (20 mg, 40 μmol) was dissolved in DMF (400 μL). TEA (19 μL, 120 μmol) was added and the solution was cooled to 0 °C. TSTU (13 mg, 44 μmol) was added and the reaction mixture was stirred for 2 h while slowly warming to room temperature. It was then concentrated under reduced pressure onto silica gel (100 mg) and the product was purified by flash chromatography on silica gel eluting with 10% MeOH in DCM. Compound **8** was obtained in 50% yield (12 mg) as a white solid. ^1^H NMR (500 MHz, DMSO-d6) δ 7.79 (s, 1H), 7.45 (d, *J* = 7 Hz, 2H), 7.31 (d, *J* = 7 Hz, 2H), 4.90 (s, 2H), 4.15-4.36 (m, 2H), 3.12-3.50 (m, 5 H), 2.71-2.98 (m, 2H), 2.51 (d, *J* = 7H, 4H), 2.00 (t, *J* = 7 Hz, 2H), 1.56-1.64 (m, 6H). (133 MHz, DMSO-d6) δ 177.9, 166.3, 164.9, 164.3, 156.1, 139.0, 128.7, 128.1, 127.7, 121.5, 67.8, 62.5, 62.2, 57.5, 38.7, 38.4, 38.1, 37.0, 27.0, 24.3, 21.8, 21.1.
